# Ambiguous support for extra-tropical accelerated diversification of rosids

**DOI:** 10.1101/2020.12.07.411439

**Authors:** Alexander Gamisch

## Abstract

Sun et al.^1^ used a comprehensive phylogenetic and a locality based climatic dataset to examine how past climates have driven diversification across 17 orders of rosids. They concluded that tropical rosids diversify slower than the (younger) non-tropical counterparts in part due to a strong negative relationship between paleo-temperature and diversification (i.e. higher diversification rates under lower temperatures). Their conclusions are based on tip rates (derived from BAMM^2^; DR^3^) and correlations with current temperature (e-SIM^4^) or binary tropicality data sets (BAMM/STRAPP^5^; FiSSE^6^) as well as tree-wide estimates of diversification with paleo-temperature (BAMM; RPANDA^7^) or tropicality (BiSSE^8^; HiSSE^9^). Here, I highlight several inconsistencies in their diversification analyses as well as a systematic error pertaining to the RPANDA model selection procedure, which, together with several minor technical issues, weaken the support for Sun et al.’s^1^ conclusions. A re-analysis of their BiSSE/HiSSE and RPANDA analyses are performed.

## Comment

### HiSSE issue

The authors assessed the relationship between tropicality/non-tropicality and diversification under a model comparison framework using four alternative BiSSE/HiSSE models. According to the authors, under both definitions of tropicality (Köppen-Geiger and geographic) the full HiSSE model fitted best, which would indicate that higher diversification of non-tropical species is modulated by an unobserved trait (Sun et al.^1^ and their Supplemental Table 4). However, when the analyses are repeated by running the authors’ code^10^ a different picture emerges (Table 1). Only under the geographic definition of tropicality the authors’ favoured model (HiSSE_full) is supported (Table 1) but with only a slightly higher net diversification rate of non-tropical species (*r*_non-tropicalA_ = 0.12 vs. *r*_tropicalB_ = 0.08). In contrast, under the Köppen-Geiger classification, a character-independent model (HiSSE_cid2_null) is favoured (Table 1). In consequence, support for (hidden) tree-wide differential character dependent diversification of tropical and non-tropical rosids is ambiguous at best.

**Table 1.** Re-run of the BiSSE/HiSSE analyses for investigating correlation between rosid diversification and climatic and geographic tropicality temperature traits using the code10 provided by the authors. Note, the HiSSE_full model is supported only for the geographic dataset in contrast to Sun et al.^1^ (cf. their Supplemental Table S4). λ, speciation rate; μ, extinction rate; r, net diversification rate (λ-μ); N, non-tropical; T, tropical; A, hidden state A; B, hidden state B.

| Traits | Model | lnL | AICc | ΔAICc | λN | μN | rN | λT | μT | rT | λNA | μNA | rNA | λTA | μTA | rTA | λNB | μNB | rNB | λTB | μTB | rTB |
| --- | --- | --- | --- | --- | --- | --- | --- | --- | --- | --- | --- | --- | --- | --- | --- | --- | --- | --- | --- | --- | --- | --- |
| Köp | BiSSE_full | -60254.92 | 120517.8 | 5476.1 | 0.45 | 0.40 | 0.06 | 0.45 | 0.39 | 0.05 | - | - | - | - | - | - | - | - | - | - | - | - |
|  | BiSSE_null | -60255.65 | 120517.3 | 5475.6 | 0.45 | 0.39 | 0.05 | 0.45 | 0.39 | 0.05 | - | - | - | - | - | - | - | - | - | - | - | - |
|  | HiSSE_full | -57643.09 | 115298.2 | 256.5 | - | - | - | - | - | - | 0.02 | 0.02 | 0.004 | 0.03 | 0.02 | 0.005 | 0.78 | 0.65 | 0.13 | 0.67 | 0.56 | 0.11 |
|  | **HiSSE_cid2_null** | **-57516.87** | **115041.7** | **0** | - | - | - | - | - | - | **0.03** | **0.03** | **0.003** | **0.03** | **0.03** | **0.003** | **0.89** | **0.80** | **0.08** | **0.89** | **0.80** | **0.08** |
| Geo | BiSSE_full | -61692.79 | 123393.6 | 5094.5 | 0.57 | 0.50 | 0.07 | 0.41 | 0.36 | 0.05 | - | - | - | - | - | - | - | - | - | - | - | - |
|  | BiSSE_null | -61886.74 | 123779.5 | 5480.4 | 0.47 | 0.42 | 0.06 | 0.47 | 0.42 | 0.06 | - | - | - | - | - | - | - | - | - | - | - | - |
|  | **HiSSE_full** | **-59143.57** | **118299.1** | **0** | - | - | - | - | - | - | **1.03** | **0.91** | **0.12** | **0.03** | **0.02** | **0.003** | **0.06** | **0.05** | **0.01** | **0.69** | **0.61** | **0.08** |
|  | HiSSE_cid2_null | -59798.55 | 119605.1 | 1306 | - | - | - | - | - | - | 0.68 | 0.45 | 0.23 | 0.68 | 0.45 | 0.23 | 0.14 | 0.09 | 0.05 | 0.14 | 0.09 | 0.05 |

### Tip rates, sample size and *p* value issue

Sun et al.^1^ found a significant relationship between current temperature and tip speciation rates (e-SIM, *p* = 0.0007) as well as significantly different and about two times higher tip speciation rates of non-tropical vs. tropical rosid species across all rosids (FiSSE, *p* = 0). It is puzzling however why these result did not hold across most individual orders (i.e. 16 out of 17), when analysed in isolation^1^. This probably hints at a general statistical problem, that the sample size has a direct impact on the *p*-value. In other words, given a large enough sample size any effect or degree of association will be statistically significant, no matter how tiny or unimportant^11,12,13^. Consequently it seems likely that the significant results of the full tree are probably only an artefact of its superior sample size (n = 19740 tips) compared to those of its composing clades/orders (n = 5 to 5678). In support of this view, the BAMM analyses across the 17 orders of Sun et al.^1^ do not support differential diversification rates of non-tropical vs. tropical rosid species. When visually inspecting the net diversification rate-through-time plots (Sun et al.^1^ and their Fig. 3) it appears that tropical and non-tropical species most often only show slight differences in net rate towards the present (viz. tips). Therefore, it is not surprising that the STRAPP analyses also do not find significant differences in tip rates *between* tropical and non-tropical species (Sun et al.^1^ and their Supplemental Table 3) (with the exception of Fabales). That being said, admittedly there seems to be geographic structure of tip rates with highest rates of diversification at northerly latitudes (e.g. in Canada, Russia) where species also tend to be youngest (c. < 1.5 million years; Sun et al.^1^ and their Fig. 2). This indicates that those northern species likely disproportionally profited from historic climate change during the Pleistocene (cf. Sun et al.^1^). Yet the number of species with high speciation rate at northern latitudes (c. 730 species with MAT < −5°C) is probably too low to drive differences between tropical and non-tropical species as a whole.

### Sensitivity test issue

It remains unclear what the STRAPP sensitivity test is supposed to achieve: for instance, if the rates between tropical and non-tropical species were not significantly different in the first place^1^, what is expected from the reduced data set? Further, the sensitivity test was done only for STRAPP permutations but without rerunning the underlying BAMM analyses (cf. Igea & Tanentzap^14^). Hence, the effect of sampling differences (i.e. the underrepresentation of tropical species^1,14^) on the upstream estimated BAMM rates per se (as well as on rate estimates derived from the other methods i.e. FiSSE, BiSSE/HiSSE, e-SIM) remains unexplored.

### RPANDA issue A

Sun et al.^1^ identified a strong negative relationship between paleo-temperature and rosid diversification (i.e. higher diversification rates under lower temperatures). This conclusion hinges on the correlation between BAMM derived net diversification rates through time across 17 orders of rosids and paleo-temperature. However visual inspection of the rate trough time plots (Sun et al.^1^ and their Fig. 3) indicate that tropical and non-tropical species most often follow near identical rate trajectories through time indicating that also historically rates of tropical and non-tropical species have never been dramatically different. This raises the question, if the diversification process would (negatively) depend on temperature would we not see historical differences in net rate between tropical and non-tropical species? Further, because young clades often show higher diversification rates^15,16^ and because temperature and age are themselves strongly auto-correlated (R^2^ = 0.78; slope = 0.21; *P* < 0.001) (i.e. temperature is decreasing towards the present) an additional independent analyses is needed^1^ to tease apart effects of temperature from those of age on diversification. For this purpose, the authors tested the fit of 18 alternative diversification models (nine time-dependent and nine temperature dependent models) for each of the 17 rosid orders within the RPANDA framework^7^. The authors found consistent support across each of the 17 orders for a model with constant speciation and extinction with respect to temperature (bcst.dcst.x)^1^.

Here, I demonstrate that the best fitting temperature-dependent model (bcst.dcst.x)^1^ was selected due to a systematic error of the RPANDA model selection procedure. In my opinion, this model definition (constant rates of λ and μ with respect to temperature) is conceptually challenging because in general rates can either vary with time or an environmental variable (e.g. temperature) or be constant. They cannot be constant with temperature, because constant rates are by definition independent from time or an environmental variable. This oxymoron applies to six out of nine temperature-dependent-diversification models (i.e. bcst.d0.x, bcst.dcst.x, bvar.dcst.x, bvar.l.dcst.x, bcst.dvar.x, bcst.dvar.l.x; see also Supplemental Method 5 of Sun et al.^1^ for model details). In fact, when looking up the corresponding code for the RPANDA analyses^10^ it turned out that for all models with a proclaimed “constant” dependency between temperature and λ and/or μ, in fact, a linear dependency (in *f.lamb* and/or *f.mu*) has been specified (e.g. *f.lamb = function(t,x,y){y[1]*x*} instead of *f.lamb = function(t,x,y){y[1]}*, where t is time and x is temperature). In other words, the authors specified a linear function in which λ and/or μ depend(s) on temperature (and therefore the rates are *not* constant), whereas the change factor (α) is apparently identical with λ at temperature 0 °C (cf. Condamine et al.^17^). Whether this special scenario is biologically plausible is open for debate; however, more importantly in the present context, the authors specified λ and/or μ to be constant (e.g. *cst.lamb =TRUE*). This latter setting simplifies the computation and should only be done when λ and/or μ are not dependent on time or the environmental variable, as explicitly stated in the RPANDA documentation^18^. These conflicting settings (specifying a variable function but implying constant rates) has an important influence on the authors’ model fitting procedure, leading to erroneous selection of the best fitting temperature-dependent model (see below).

### RPANDA issue B

I noted an additionally error on how incomplete taxon sampling was accounted for in the RPANDA analyses. Looking at the code^10^, it appears that the sampling fraction of each clade was taken as fraction of the total tree (and not as the fraction of sampled species versus the total number of clade species; cf. the BAMM analyses of Sun et al.^1^). For example, the Geraniales clade has the highest relative sampling fraction of all orders (305 sampled out of total of 967 species; sampling fraction = 0.315; see Supplemental Table 7 of Sun et al.^1^); however, according to the RPANDA code provided by the authors^10^ (i.e. *fraction <-length(tree$tip.label)/NN*; whereas NN is the length of the full rosid tree), the sampling fraction would be 0.015 (305 out of 19,740 species). Since the various likelihood expressions depend on the fraction of species that are represented in the phylogeny^7,19^ the deflated sampling fractions are likely to have been further detrimental to the accuracy of the RPANDA parameter estimates and model selection procedure (e.g. Sun et al.^20^).

### RPANDA issue C

A final issue relates to the recent criticism of linear diversity dependencies in RPANDA^21^, which is therefore relevant not only to the study of Sun et al.^1^ but also to similar ones using this R package. In short, linear diversification dependencies can lead to negative λ and/or μ rates that would be positivised in the current implementation of RPANDA^21,22^. In such cases, a rate curve is fitted that is first decreasing backwards in time and then increasing again (e.g. Supplementary Fig. 1), leading to biologically questionable scenarios^21,22^. When performing initial test runs, I identified that this linear diversity-dependency issue in RPANDA^21^ also applies to some of the models tested by the authors (e.g. bvar.l.dcst.x, bcst.dvar.l.x; Supplementary Fig. 1). Specifying an alternative function (e.g. λ(t) = max(0, λ0+αt); *sensu* Morlon et al.^22^) did not work in these instances; therefore, I excluded all linear (time and temperature) dependencies from the reanalyses (*sensu* Gamisch^21^) with the exception of the bcst.dcst.x model for the purpose of demonstrating RPANDA issue A (see above).

Results indicate that the conflicted (linear) temperature-dependent model with λ and μ set constant (here bcst.dcst.x_cst; Table 2) produces superior AICc scores that are about 9 to 20 % lower than the otherwise identical model for which rates are not set constant (bcst.dcst.x_cor; Table 1). This indicates that the best fitting constant speciation and extinction model with respect to temperature across the 17 orders of rosids was likely identified based on a systematic error in the model selection procedure. Moreover, when the analyses were repeated with the corrections detailed above, a more complex picture emerged with regard to the temperature-dependent diversification of rosids, i.e. across the 17 orders and the full tree, eight out of 18 clades unambiguously best fitted only a single model (based on ΔAICc > 2.0) (Table 2). Of those eight clades, six favoured time-dependent models, including constant (Crossosomatales, Huerteales, Picramniales, Zygophyllales), exponential increasing (Cucurbitales), and wax and waning (Brassicales) net diversification rates (Fig. 1). The remaining two clades (Fagales and Geraniales) favoured temperature-dependent diversification, whereby the relationship between temperature and diversification was negative in Fagales (r increasing with lower temperature towards the present) but positive in Geraniales (r decreasing with lower temperature towards the present). The remaining nine clades plus the full tree favoured multiple models with essentially equivalent fit (Table 2). Such ambiguous support for multiple different models may arise due to the consistence of multiple diversification histories with a given phylogeny^23^ or simply the different processes that may drive diversification across different parts of a (large) phylogeny. Nevertheless, when considering the best temperature-dependent model among the remaining ten models with ambiguous model support (Table 1, Fig. 1), decreasing r with lower temperature towards the present was found in eight out of ten clades (Celastrales, Fabales, Malpighiales, Malvales, Myrtales, Oxalidales, Sapindales, full tree). Only two orders (Rosales, Vitales) showed ambiguous support for a negative relationship between temperature and diversification (Table 2, Fig. 1).

**Table 2.**
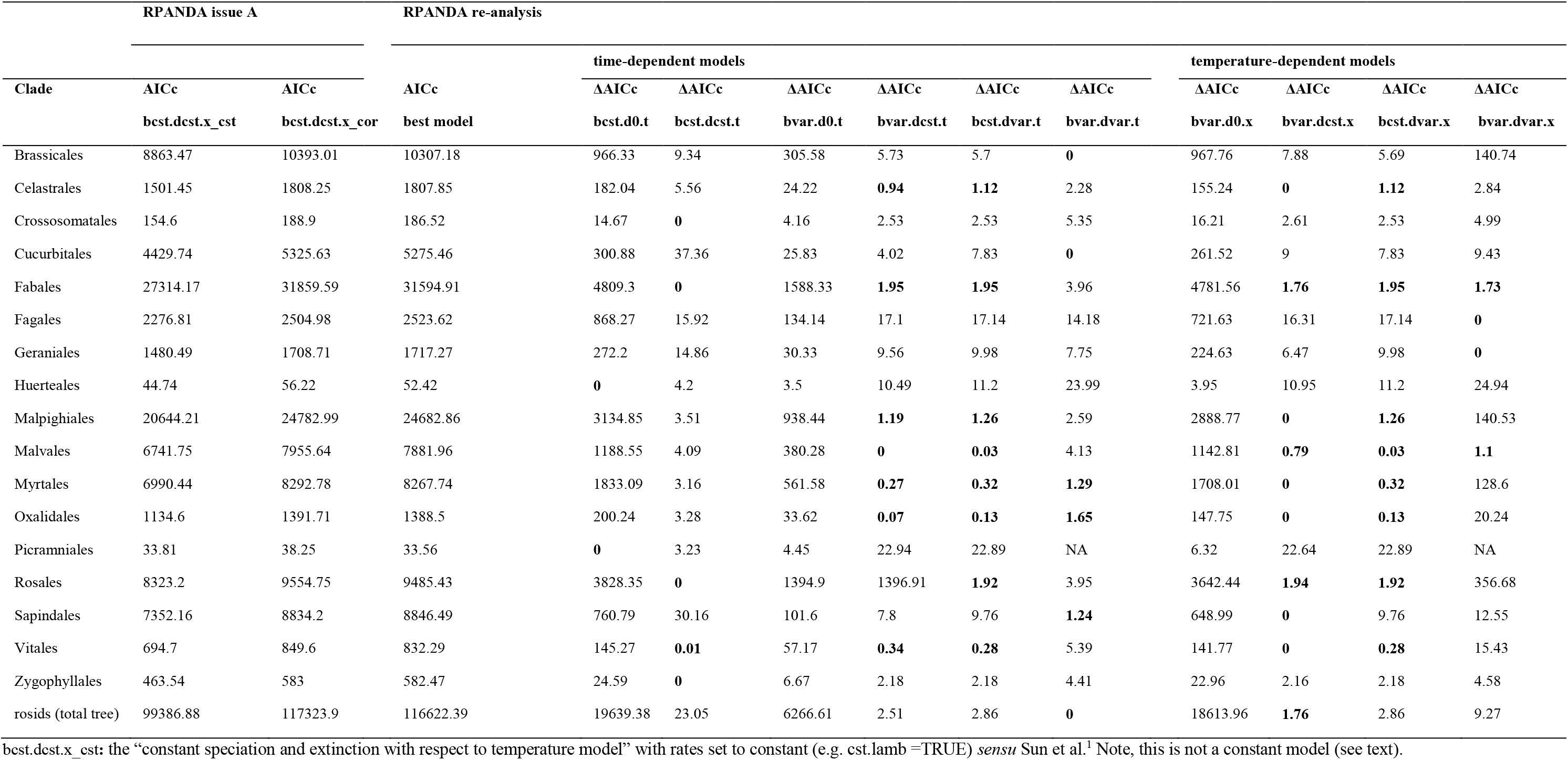
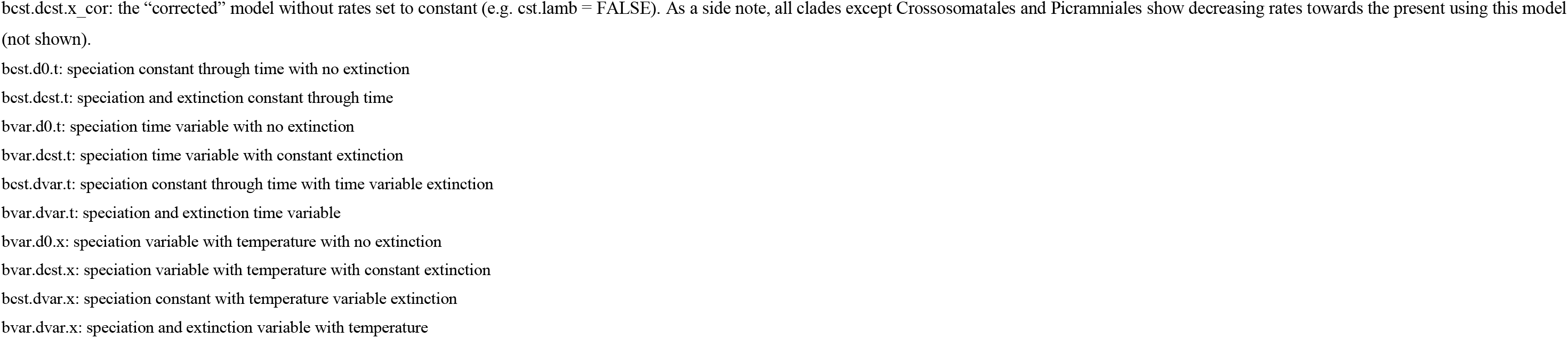
Demonstration of the systematic error (RPANDA issue A) and RPANDA reanalysis. Model support for ten (six time-dependent and four temperature-dependent) models of diversification across each of the 17 rosid orders as well as the full tree, using RPANDA and taking clade specific sampling fractions into account. With the exception of RPANDA issue A, only exponential relationships with time or temperature have been considered (see text). The models with ΔAICc < 2.0 are indicated in bold. See footnotes for explanation of model acronyms.

**Fig. 1.**
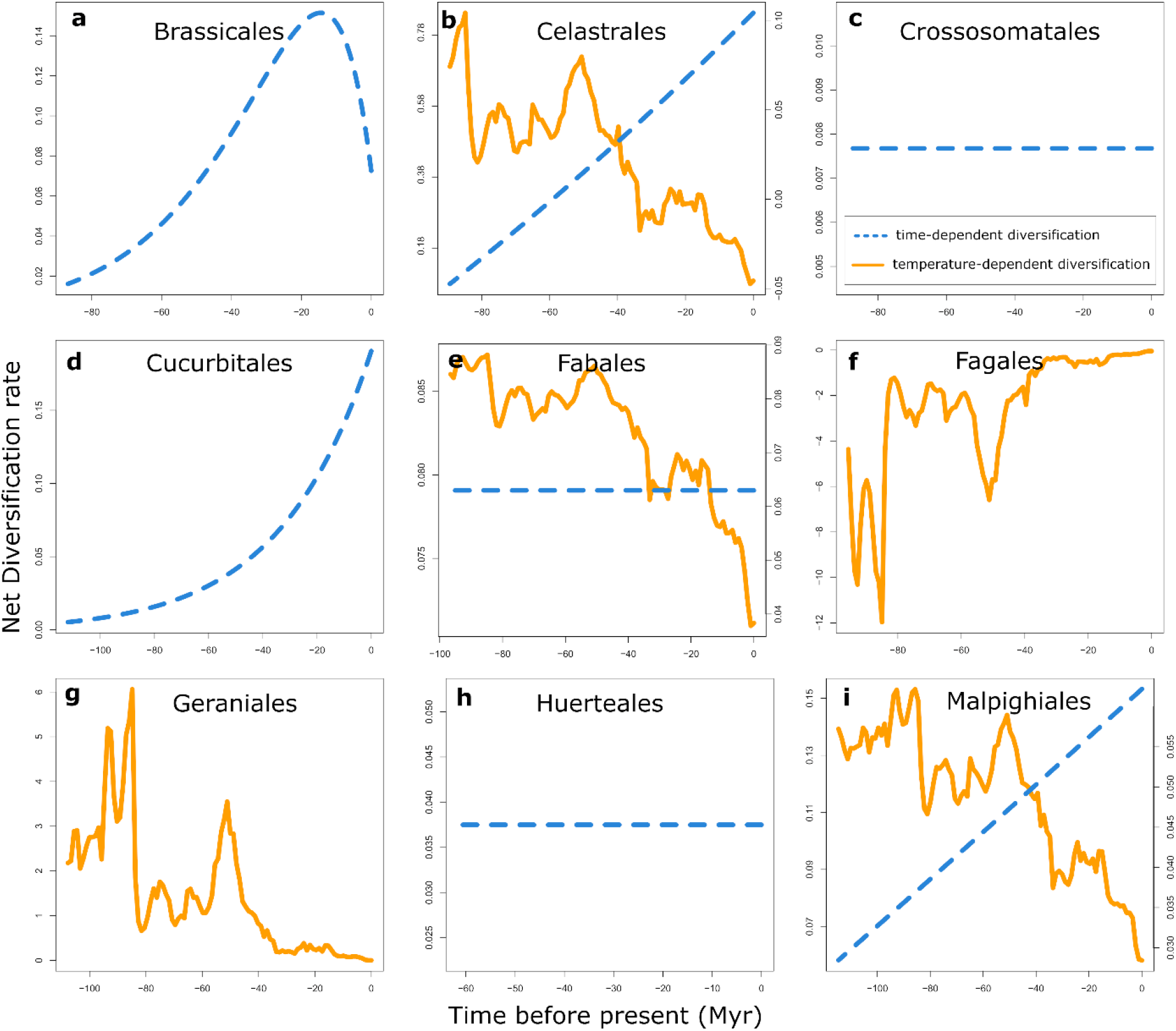
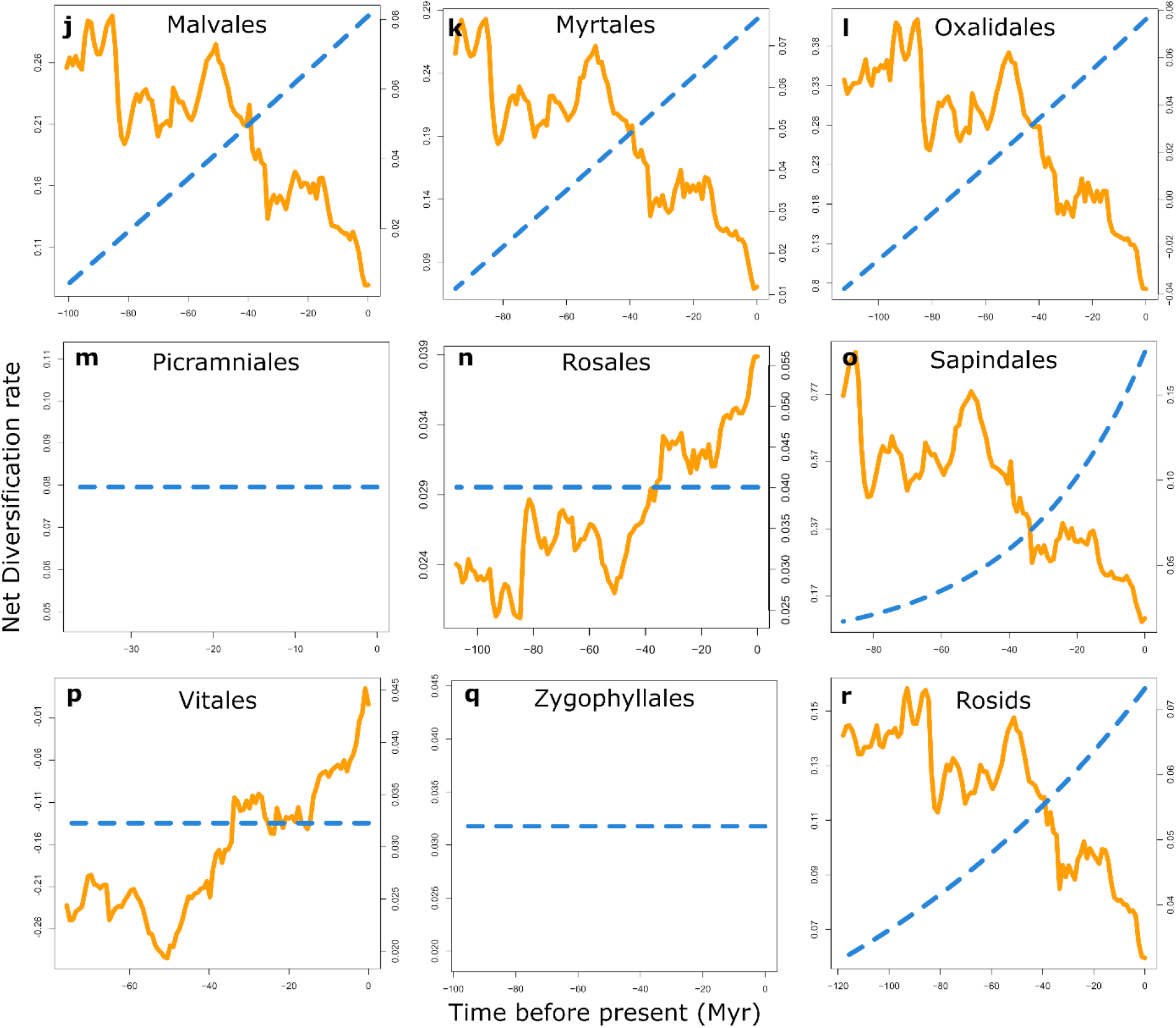
Net diversification rate-through-time plots of the best fitting models of time- and/or temperature-dependent diversification for each of the 17 rosid orders (clades) and the full tree (**a-r**) as estimated with RPANDA. Graphs show either the best fitting models (based on ΔAICc > 2.0) or the best time-dependent and temperature-dependent models when they had essentially equivalent fit (< 2 ΔAICc units). See also Table 2 for details. Myr, million years.

In summary, I highlight several issues with the diversification analyses of Sun et al.^1^ I could only partly reproduce the results of their HiSSE analyses leading to ambiguous support for they favoured model. Further, I stress that the significant association between tip rates and temperature or tropicality are probably an artefact of sample size. Both tropical and non-tropical rosids, like many other plant lineages (e.g. Gamisch & Comes^24^; Huang et al.^25^), apparently showed accelerated diversification from the mid-Miocene onwards (within the last 10–15 million years; cf. Sun et al.^1^). The present RPANDA re-analyses also indicate that the diversification of rosids can probably not be explained by a negative paleo-temperature-dependence of the diversification process. Instead, for most clades (and the full rosid tree) there is at least ambiguous support for an alternative explanation of higher tropical species richness (i.e. initially higher rates under higher temperatures; e.g. Mittelbach et al.^26^, Rolland et al.^27^, Shrestha et al.^28^ and references therein). Thus, notwithstanding that young northern latitude species tend to have high rates^1^, there seems to be no strong evidence that cooling per se had a disproportionate effect on non-tropical rosid diversification.

## Methods

The BiSSE/HiSSE analyses were repeated by running the authors’ HiSSE code^10^ using hisse v1.9.8^9^. To demonstrate RPANDA issue A, I used RPANDA v1.7 and the authors’ RPANDA code^10^ (see Supplemental Material) to rerun their best fitting model with λ and μ set constant (e.g. cst.lamb =TRUE; like the authors did; here bcst.dcst.x_cst) and without this setting (e.g. cst.lamb =FALSE; bcst.dcst.x_cor). For the RPANDA reanalyses, I fitted ten non-linear models of diversification across each of the 17 rosid orders (clades) as well as the full tree (i.e. six time-dependent and four temperature-dependent models; see Table 2) following the authors’ RPANDA code^10^ (see Supplemental Material) but with clade specific sampling fractions taken into account (see Sun et al.^1^ and their Supplemental Table 7). Due to the RPANDA linear diversity-dependency issue^21^ only exponential relationships with time or temperature have been considered. Net diversification rate-trough-time plots of the best fitting models were plotted using the *plot_fit_bd* or *plot_fit_env* functions of RPANDA. The best fitting models were selected based on ΔAICc > 2.0^29^.

## Acknowledgments

I want to thank the corresponding authors of Sun et al. for discussions as well as for providing intermediate results of their RPANDA analyses. Special thanks are expressed to Jeremy Beaulieu (University of Arkansas) for fruitful discussions about the HiSSE 32 vs. 64 bit issue and to Hans Peter Comes (University of Salzburg) for very helpful comments on earlier versions of this manuscript. The present work was funded by the FWF (Austrian Science Fund) grant P29371 to Hans Peter Comes.

## Author Contributions

AG designed the study, has analysed the data and has written the manuscript.

## Data Availability

All data used in this study is available from Sun and Folk (2020)^10^.

## Code availability

The code used for the demonstration of RPANDA issue A and the RPANDA reanalysis is available as Supplementary Material.

