## Supplementary Material for "Ambiguous support for extra-tropical accelerated diversification of rosids"

Supplementary Information:
Supplementary Figure: 1
Supplementary Method (R Code)

**Supplementary Figure**

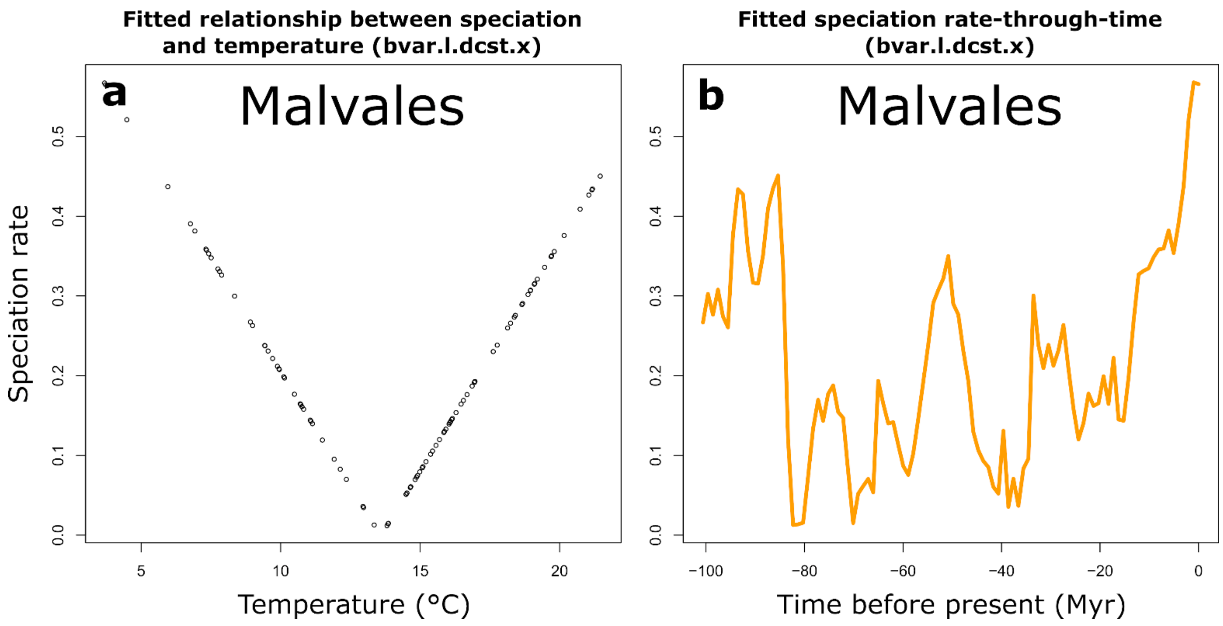


**Supplementary Fig. 1.** Demonstration of the effect of the linear diversification dependency issue (RPANDA issue C;) on the fitted relationship between speciation and temperature using the using the Malvales clade and a linear temperature dependent model (speciation variable with temperature with constant extinction; bvar.l.dcst.x) as an example. In a) speciation rate is highes at lowest temperature and then speciation rate decreases until c. 14 °C from where it increases again with high rates at > 20°C. This increase from 14 °C happens because the rate would be negative according the linear function but RPANDA (fit_bd, fit_env) uses absolute values (see Gamisch[21] for details). b) As a consequence speciation rates are modelled as high early and late in the clades history when global temperature was high and low, respectively.

**Supplementary Method**

R code:

```
#### This code is built upon the datasets and code provided by Sun et al. #####
####(https://doi.org/10.5281/zenodo.3843441)
#### and is used to demonstrate the RPANDA issue A as well as to perform the RPANDA
#### reanalysis.
#### NOTE: Here the analyses is done not in batch but on a tree by tree basis (the models are
#### run for each tree seperately).

##SETUP

# download the datasets of Sun et al. (https://doi.org/10.5281/zenodo.3843441)
# copy the trees (.tre) and paleotemperature file (Global_paleo-temperature.csv) into a single
# folder and select as working directory

#loading packages
library("RPANDA")
library("ape")
library("phytools")

# load one a of the trees
tree <-read.tree("Rosales_5g.tre")
tree <-read.tree("Zygophyllales_5g.tre")
tree <-read.tree("Vitales_5g.tre")
tree <-read.tree("Sapindales_5g.tre")
tree <-read.tree("Picramniales_5g.tre")
tree <-read.tree("Oxalidales_5g.tre")
tree <-read.tree("Myrtales_5g.tre")
tree <-read.tree("Malvales_5g.tre")
tree <-read.tree("Malpighiales_5g.tre")
tree <-read.tree("Huerteales_5g.tre")
tree <-read.tree("Geraniales_5g.tre")
tree <-read.tree("Fagales_5g.tre")
tree <-read.tree("Fabales_5g.tre")
tree <-read.tree("Cucurbitales_5g.tre")
tree <-read.tree("Crossosomatales_5g.tre")
tree <-read.tree("Brassicales_5g.tre")
tree <-read.tree("Celastrales_5g.tre")
tree <-read.tree("rosids_5g_whole_tree.tre")

# specify clade specific sampling fractions for the tree selected above
fraction <-     0.308   # Brassicales
fraction <-     0.187   # Celastrales
fraction <-     0.293   # Crossosomatales
fraction <-     0.278   # Cucurbitales
fraction <-     0.235   # Fabales
fraction <-     0.237   # Fagales
fraction <-     0.315   # Geraniales
fraction <-     0.233   # Huerteales
```

```
fraction <-      0.18     # Malpighiales
fraction <-      0.173   # Malvales
fraction <-      0.088   # Myrtales
fraction <-      0.087   # Oxalidales
fraction <-      0.088   # Picramniales
fraction <-      0.086   # Rosales
fraction <-      0.193   # Sapindales
fraction <-      0.103   # Vitales
fraction <-      0.206   # Zygophyllales
fraction <-      0.172   # rosids_whole_tree

#loading temperature data
Tem.C <- read.csv("Global_paleo-temperature.csv", as.is = TRUE)
dof<-smooth.spline(Tem.C[,1], Tem.C[,2])$df

# caculate crown age
tot_time<-max(node.age(tree)$ages)

# define initial values
 par <- list(c(0.09), c(0.09, 0.005), c(0.05, 0.01), c(0.05, 0.01), c(0.05, 0.01, 0.005), c(0.09,
0.001, 0.005), c(0.05, 0.005, 0.01),
            c(0.05, 0.005, 0.001), c(0.05, 0.01, 0.005, 0.0001))

## now everthing is ready to run the models below

#######RPANDA_Models#############

#### RPANDA ISSUE A

  #11)   Constant speciation and extinction with temperature (x) (here as bcst.dcst.x_cst)
  f.lamb.x = function(t,x,y){y[1]*x}
  f.mu.x = function(t,x,y){y[1]*x}

  model <- "bcst.dcst.x_cst"
  bcst.dcst.x_cst <- fit_env(tree, Tem.C, tot_time, f.lamb.x, f.mu.x, lamb_par=par[[2]][1],
mu_par=par[[2]][2], cst.lamb=TRUE, cst.mu=TRUE, cond="crown", f=fraction, df=dof,
dt=1e-3)

  #11cor)         Constant speciation and extinction with temperature (x) (here as
bcst.dcst.x_cor)
  f.lamb.x = function(t,x,y){y[1]*x}
  f.mu.x = function(t,x,y){y[1]*x}

  model <- "bcst.dcst.x_cor"
  bcst.dcst.x_cor <- fit_env(tree, Tem.C, tot_time, f.lamb.x, f.mu.x, lamb_par=par[[2]][1],
mu_par=par[[2]][2], cond="crown", f=fraction, df=dof, dt=1e-3)

# print and inspect model results

print(bcst.dcst.x_cst)
print(bcst.dcst.x_cor)
```

```
# plot the diversification rates through time of the temperature dependent models

plot_fit_env(bcst.dcst.x_cor,Tem.C,tot_time)

#### RPANDA REANALYSIS

#Six time-dependent models (note the model numbers refer to the original nummeration of Sun et al.)

  #1)    Pure birth model, no extinction rate (mu, μ = 0), and constant speciation rate (lambda, λ; hereafter bcst.d0.t)
  # t is time
  # y is a vector of initial values feeding to the functions of λ and μ
  f.lamb.t = function(t, y){y[1]}
  f.mu.t = function(t, y){0}

  model <- "bcst.d0.t"
  bcst.d0.t <- fit_bd(tree, tot_time, f.lamb.t, f.mu.t, lamb_par=par[[1]][1], mu_par=c(), f=fraction, cst.lamb=TRUE, fix.mu=TRUE, cond="crown", dt=1e-3)

  #2)    Birth-death model with constant speciation and extinction (here as bcst.dcst.t)
  f.lamb.t = function(t, y){y[1]}
  f.mu.t = function(t, y){y[1]}

  model <- "bcst.dcst.t"
  bcst.dcst.t <- fit_bd(tree, tot_time, f.lamb.t, f.mu.t, lamb_par=par[[2]][1], mu_par=par[[2]][2], cst.lamb=TRUE, cst.mu=TRUE, cond="crown", f=fraction, dt=1e-3)

  #3)    Pure birth model with exponential variation in speciation rate (here as bvar.d0.t)
  f.lamb.t = function(t, y){y[1] * exp(y[2] * t)}
  f.mu.t = function(t, y){0}

  model <- "bvar.d0.t"
  bvar.d0.t <- fit_bd(tree, tot_time, f.lamb.t, f.mu.t, lamb_par=par[[3]][c(1,2)], mu_par=c(), expo.lamb=TRUE, fix.mu=TRUE, cond="crown", f=fraction, dt=1e-3)

  #5)    Birth-death model with exponential variation in speciation rate and constant extinction (here as bvar.dcst.t)
  f.lamb.t = function(t, y){y[1] * exp(y[2] * t)}
  f.mu.t = function(t, y){y[1]}

  model <- "bvar.dcst.t"
  bvar.dcst.t <- fit_bd(tree, tot_time, f.lamb.t, f.mu.t, lamb_par=par[[5]][c(1,2)], mu_par=par[[5]][3], expo.lamb=TRUE, cst.mu=TRUE,cond="crown", f=fraction, dt=1e-3)

  #7)    Birth-death model with a constant speciation rate and exponential variation in extinction (here as bcst.dvar.t)
  f.lamb.t = function(t, y){y[1]}
  f.mu.t = function(t,y){y[1] * exp(y[2] * t)}
```

```
  model <- "bcst.dvar.t"
  bcst.dvar.t <- fit_bd(tree, tot_time, f.lamb.t, f.mu.t, lamb_par=par[[7]][1],
mu_par=par[[7]][c(2,3)], cst.lamb=TRUE, expo.mu=TRUE, cond="crown", f=fraction,
dt=1e-3)

  #9)    Birth-death model with exponential variation in speciation and extinction (here as
bvar.dvar.t)
  f.lamb.t = function(t, y){y[1] * exp(y[2] * t)}
  f.mu.t = function(t,y){y[1] * exp(y[2] * t)}

  model <- "bvar.dvar.t"
  bvar.dvar.t <- fit_bd(tree, tot_time, f.lamb.t, f.mu.t, lamb_par=par[[9]][c(1,2)],
mu_par=par[[9]][c(3,4)], expo.lamb=TRUE, expo.mu=TRUE, cond="crown", f=fraction,
dt=1e-3)

#Four temperature-dependent models

  #12)   Exponential variation in speciation rate with temperature (x) (here as bvar.d0.x)
  f.lamb.x = function(t,x,y){y[1] * exp( y[2] * x)}
  f.mu.x = function(t,x,y){0}

  model <- "bvar.d0.x"
  bvar.d0.x <- fit_env(tree, Tem.C, tot_time, f.lamb.x, f.mu.x, lamb_par=par[[3]][c(1,2)],
mu_par=c(), expo.lamb=TRUE, fix.mu=TRUE, cond="crown", f=fraction, df=dof, dt=1e-3)

  #14)    Exponential variation in speciation rate with temperature (x) and constant extinction
(here as bvar.dcst.x)
  f.lamb.x = function(t,x,y){y[1] * exp( y[2] * x)}
  f.mu.x = function(t,x,y){y[1]}

  model <- "bvar.dcst.x"
  bvar.dcst.x <- fit_env(tree, Tem.C, tot_time, f.lamb.x, f.mu.x, lamb_par=par[[5]][c(1,2)],
mu_par=par[[5]][3], expo.lamb=TRUE, cst.mu=TRUE,cond="crown", f=fraction, df=dof,
dt=1e-3)

  #16)   Constant speciation and exponential variation in extinction with temperature (x) (here
as bcst.dvar.x)
  f.lamb.x = function(t,x,y){y[1]}
  f.mu.x = function(t,x,y){y[1] * exp(y[2] * x)}

  model <- "bcst.dvar.x"
  bcst.dvar.x <- fit_env(tree, Tem.C, tot_time, f.lamb.x, f.mu.x, lamb_par=par[[7]][1],
mu_par=par[[7]][c(2,3)], cst.lamb=TRUE, expo.mu=TRUE, cond="crown", f=fraction,
df=dof, dt=1e-3)

  #18)   Exponential variation both in speciation and extinction rates with temperature (x) (here
as bvar.dvar.x)
  f.lamb.x = function(t,x,y){y[1] * exp( y[2] * x)}
  f.mu.x = function(t,x,y){y[1] * exp(y[2] * x)}

  model <- "bvar.dvar.x"
```

```
  bvar.dvar.x <- fit_env(tree, Tem.C, tot_time, f.lamb.x, f.mu.x, lamb_par=par[[9]][c(1,2)],
mu_par=par[[9]][c(3,4)], expo.lamb=TRUE, expo.mu=TRUE, cond="crown", f=fraction,
df=dof, dt=1e-3)

###############

# print and inspect model results

print(bcst.d0.t)
print(bcst.dcst.t)
print(bvar.d0.t)
print(bvar.dcst.t)
print(bcst.dvar.t)
print(bvar.dvar.t)
print(bvar.d0.x)
print(bvar.dcst.x)
print(bcst.dvar.x)
print(bvar.dvar.x)


# plot the diversification rates through time of the time dependent models, e.g.:

plot_fit_bd(bvar.dvar.t,tot_time)

# plot the diversification rates through time of the temperature dependent models, e.g.:

plot_fit_env(bvar.dvar.x,Tem.C,tot_time)
```
